# A Plug-and-Play P300-Based BCI with Calibration-Free Application

**DOI:** 10.1101/2025.05.21.655021

**Authors:** Jongsu Kim, Sung-Phil Kim

## Abstract

The practical deployment of P300-based brain–computer interfaces (BCIs) have long been hindered by the need for user-specific calibration and multiple stimulus repetitions. In this study, we build and validate a plug-and-play, calibration-free P300 BCI system that operates in a single-trial setting using a pre-trained xDAWN spatial filter and a deep convolutional neural network. Without any subject-specific adaptation, participants could control an IoT device via the BCI system in real time with decoding accuracy reaching 85.2% comparable to the offline benchmark of 87.8%, demonstrating the feasibility of realizing a plug-and-play BCI. Offline analyses revealed that a small set of parietal and occipital electrodes contributed most to decoding performance, supporting the viability of low-density, high-accuracy BCI configurations. A data sufficiency simulation provided quantitative guidelines for pre-training dataset size, and an error trial analysis showed that both stimulus timing and preparatory attentional state influenced real-time decoding performance. Together, these results demonstrate the first real-time validation of a calibration-free, single-trial P300 BCI and offer practical insights for developing scalable, robust, and user-friendly BCI systems.

## I. Introduction

**B**rain-computer interfaces (BCIS) are a rapidly growing technology that enable users to interact with computers, machines, or digital environments directly through brain activity, without the need for any physical movement or muscular control [1]. BCIs interpret neurophysiological signals to translate the user’s intention into actionable commands. They hold tremendous promise across a wide range of applications—from assistive technologies for individuals with severe motor impairments, to hands-free control of smart devices, neurorehabilitation systems, and gaming [2–5]. Among the three main paradigms of BCIs—active, reactive, and passive—reactive BCIs have received significant attention due to their high reliability and relatively low training burden [6, 7]. In reactive BCI paradigms, users respond passively to external stimuli, and their brain’s involuntary responses are decoded to infer intent.

Two of the most established paradigms in reactive BCI research utilize steady-state visual evoked potentials (SSVEP) [8] and P300-based event-related potentials (ERPs) [9]. SSVEP-based BCIs detect periodic EEG responses that are time-locked to the frequency of a flickering visual stimulus. These responses are strong, consistent, and relatively easy to classify, which enables SSVEP systems to support high-speed control. However, SSVEP systems require specific technical conditions such as high-refresh-rate monitors to create stable flickering and their performance can degrade in interfaces where visual stimuli are spatially constrained or placed too close together [10]. On the other hand, P300-based BCIs rely on detecting the P300 ERP component, a positive EEG deflection that typically occurs about 300 milliseconds after the user perceives a rare but meaningful stimulus in an “oddball” paradigm [11, 12]. Because they do not require flickering and are more flexible in terms of stimulus design, P300 BCIs can be integrated into more natural and user-friendly interfaces. They are especially suitable for applications involving discrete selection tasks, such as spelling systems, menu navigation, or command selection.

Despite their advantages, P300-based BCIs face several technical challenges that have limited their transition from laboratory settings to real-world applications. One of the primary challenges is the need for multiple trials of repeated stimulus presentations to obtain reliable ERPs [13–17]. Since single-trial P300 responses are often contaminated by noise, repeated presentations are used to average out this noise and enhance the signal-to-noise ratio [18, 19]. However, repetition significantly slows down interaction and induces user fatigue, reducing the overall efficiency of the system. Another major challenge is the reliance on user-specific calibration sessions, which are time-consuming and require users to provide labeled data before the system can be used effectively [20]. This undermines the concept of “plug-and-play” BCIs and is particularly problematic in clinical or large-scale deployment scenarios. Furthermore, there is substantial inter-subject variability in P300 responses due to differences in attention, fatigue, and cognitive engagement, which complicates the design of generalized models [21–24]. Lastly, the stimulus design itself plays a crucial role in the quality of the elicited P300. Poorly designed or low-relevance stimuli often fail to engage the user’s attention, leading to inconsistent ERP generation [25–27].

To address these issues, prior studies have explored improvements in both stimulus design and signal processing. Our previous work proposed a novel stimulus paradigm using task-relevant finger-tapping animations to simulate physical interaction, thereby enhancing attention and improving ERP responses [28]. Using this design, we demonstrated enhanced decoding performance at the single-trial level without individual calibration in an offline analysis. On the signal processing side, recent advances such as xDAWN spatial filtering [29], Riemannian geometry-based features [22], and deep learning classifiers [30] have also shown promise in improving performance across users. However, most of these findings were derived from offline evaluation in which training and testing were performed on fixed datasets with cross-validation, rather than being validated on entirely new users in real time. As such, the real-world generalizability and robustness of these approaches remain uncertain. Therefore, it remains unclear whether P300 BCI systems equipped with the novel stimulus design and advanced signal processing techniques can function robustly in real-time, user-independent settings.

In the present study, we address this critical gap by validating a calibration-free, single-trial P300 BCI system in an online experiment. Thirty-four participants interacted with a smart IoT lighting device by focusing on visually presented stimuli. The system used a pre-trained xDAWN with deep convolution network pipeline, optimized from prior offline datasets, and processed EEG signals in real time without any user-specific training. Participants issued commands such as turning the light on/off, changing color, or adjusting brightness, and the system responded instantly to their brain signals.

The results demonstrated that the proposed system achieved high classification accuracy and information transfer rate (ITR) under a fully plug-and-play setting, using only single-trial ERPs. To complement the real-time experiment, we conducted several post-hoc offline analyses to deepen our understanding of model robustness and guide future development of practical P300-based BCIs. First, a channel ablation analysis was performed to quantify the contribution of each EEG channel to classification accuracy, supporting efforts toward minimal-channel system design. Second, a data sufficiency simulation explored how many participants and blocks are needed for the training set to construct effective pre-trained decoders, providing quantitative guidelines for scalable plug-and-play P300-based BCIs. Third, an error trial analysis examined both behavioral and neural correlates of misclassifications—including alpha-band power fluctuations and stimulus sequence effects—using post-task interviews and a binomial generalized linear mixed model (GLMM).

Additionally, motivated by the observation that single-trial paradigms offer no second chance to detect missed stimuli, we explored whether a predictable (fixed-order) stimulus presentation might reduce attentional lapses and maintain performance. However, as we report in the Discussion section below, the results suggest that maintaining stimulus unpredictability remains essential even in single-trial settings. This finding underscores the importance of cognitive factors—such as attentional engagement and stimulus expectancy—that dynamically influence P300 elicitation. The on-line experimental results of single-trial, calibration-free decoding with a computationally lightweight, real-time signal processing highlight the potential of plug-and-play P300 BCIs deployable beyond controlled laboratory settings.

## II. Methods

## A. Participants

A total of fifty healthy adults participated in the online BCI experiment. Among them, thirty-four participants (mean age = 23.41 years old, SD = 4.72) completed the main experiment using a random-order stimulus presentation. Sixteen additional participants mean age = 22.44 years old, SD = 3.30) were tested under a fixed-order condition to examine whether predictable stimulus sequencing could reduce missed responses in single-trial settings. Results from the fixed-order condition are reported in the Discussion. All participants had normal or corrected-to-normal vision and no history of neurological or psychiatric disorders. Written informed consent was obtained from participants in accordance with the UNIST Institutional Review Board (UNIST-IRB-18-08-A).

### B. Experiment Design

The experiment was designed with a four-option P300 oddball task simulating smart home light control. Participants selected one of the four visual icons on a screen to issue commands to a smart light through brain signals. The four commands included turning the light on, turning it off, adjusting brightness, and changing color. Each command was represented by a distinct icon displayed at a fixed location in one of the screen’s four corners (Fig. 1A).

**Fig. 1.**
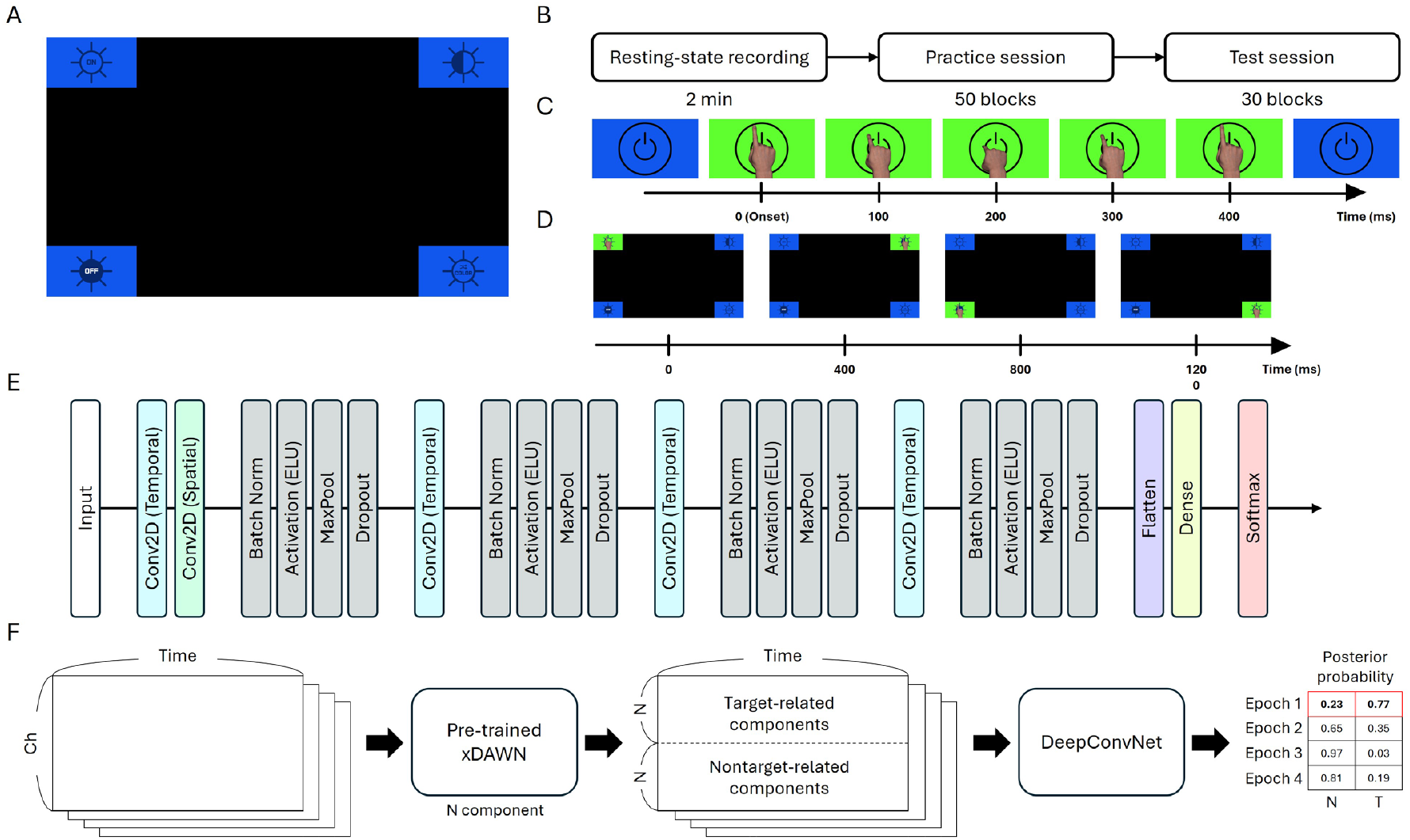
Overview of the experimental paradigm, visual stimuli, and signal processing pipeline. A) The user interface (UI) presented to participants during the P300-based IoT smart lighting control task. Four icons representing distinct light commands (e.g., power on/off, brightness adjustment, color change) were positioned at each corner of the screen. B) The overall experimental procedure, consisting of a 2-minute resting-state EEG recording, a 50-block practice session, and a 30-block test session with real-time BCI operation. C) Example of the finger-tapping animation used as visual stimuli. Each stimulus was shown for 400 ms, visually mimicking the act of pressing a virtual button. D) Illustration of stimulus presentation order in the fixed-order condition, where the four stimuli are shown sequentially in a predetermined spatial order. In the random-order condition, the same stimuli were presented in a randomized sequence (not depicted here for brevity). E) The architecture of DeepConvNet used for classifying P300 responses. The model included multiple temporal and spatial convolution layers, batch normalization, max pooling, dropout regularization, and a final softmax layer. F) The signal processing pipeline for real-time classification. EEG data were first transformed using a pre-trained xDAWN spatial filter, separating target- and non-target-related components. These components were then passed into the DeepConvNet, which computed the posterior probability for each class based on the four stimulus presentations in each trial.

To reflect realistic use, the task procedure was designed in a practical scenario. Each session began with the light turned off, and participants were first instructed to turn it on. Once the light was on, subsequent commands were randomly chosen from the remaining three options—adjusting brightness, changing color, or turning off. When the light was off, the next command always prompted the user to turn it back on. This ensured a logical cycle of interaction while maintaining variability in the task structure.

Before the main experiment, participants completed a 2-minute baseline EEG recording while fixating on a central cross and minimizing movement (Fig. 1B). The baseline was used to calibrate artifact subspace reconstruction (ASR) parameters for online preprocessing and to identify bad channels to exclude from subsequent signal processing.

Each trial consisted of the following sequence. First, a fixation cross was presented in the center of the screen for 1 second, during which participants were asked to maintain steady gaze and minimize movement. Next, one of the four icons was highlighted as the target, displayed for 1 second to indicate the command participants should focus on. This cue served as an instruction to mentally prepare for the upcoming selection process. Following the cue, the four icons were each sequentially highlighted with a 400-millisecond finger-tapping animation, which visually simulated the act of pressing the button (Fig. 1C). The four icons were highlighted once per block in a randomized order. Participants were instructed to focus on the target and ignore other non-targets. This constituted a single-trial oddball paradigm, where each option appeared once per block without repetition. A block consisted of four trials corresponding to each icon. No inter-stimulus interval was given in the block.

The experiment consisted of a practice session followed by a test session. In both sessions, participants interacted with a smart lighting device (Philips Hue; Signify N.V., Eindhoven, Netherlands) through a P300-based interface. During the practice session, system control was not based on EEG signals. Instead, light responses were automatically executed using the aforementioned scenario. Participants were informed that the system was learning from their brain signals and were instructed to concentrate as if their EEG data were actively being processed. The purpose of this approach was to encourage engagement and stabilize attentional focus, both of which are critical for eliciting reliable P300 responses. Participants were also told that automatic control of the light was intended to help them concentrate by minimizing frustration.

The practice session consisted of 50 blocks and was designed to familiarize participants with the task flow, interface, and timing of stimulus presentation. Since P300 responses are sensitive to attention and cognitive readiness, the extended number of blocks in the practice session was intended to ensure that participants reached a consistent mental state suitable for reliable ERP generation. Participants were encouraged to choose their own mental strategy—such as mental counting or imagery—rather than following a prescribed mental task.

The test session consisted of 30 blocks. In this session, EEG signals were processed and classified in real time using a pre-trained decoder (see Section II. D-F). The number of blocks was chosen to balance adequate performance assessment with minimizing fatigue, which could negatively affect attention in single-trial tasks. Based on the classification results, the system executed the corresponding light control commands, and participants received immediate visual feedback through changes in the lighting device (supplementary video 1). This allowed us to evaluate the BCI system’s real-time performance under conditions approximating practical use.

### C. EEG Data Aquisition

EEG signals were recorded using a 31-channel actiCHamp system (Brain Products GmbH, Germany) with active wet electrodes positioned according to the international 10–20 system. Electrode sites included frontal, central, parietal, and occipital regions to ensure comprehensive spatial coverage of ERP components, particularly those associated with visual attention and P300 generation. The left mastoid was used as the reference electrode and the right mastoid served as the ground. Signals were sampled at 500 Hz, and electrode impedance was kept below 10 kΩ throughout the recording. Data quality was continuously monitored to ensure minimal contamination from muscle, eye, or movement artifacts.

### D. Preprocessing and Feature Extraction

All EEG data underwent a preprocessing pipeline prior to feature extraction and classification. In both the online and offline settings, raw EEG signals were first bandpass filtered using a 1-Hz high-pass filter to remove slow drifts and a 30-Hz low-pass filter to eliminate muscle artifacts and high-frequency noise. Filters were implemented as finite impulse response (FIR) filters using a Hamming window design. Bad channels were identified based on a two-minute resting-state EEG recording acquired before the task. Following a modified version of the ‘clean_channels’ function from the EEGLAB toolbox, the correlation of each channel with its neighbors was assessed in 5-second sliding windows. Channels showing correlation coefficients below 0.8 in more than 40% of the total baseline period were flagged as bad. The same set of bad channels, identified from the baseline recording, was used throughout the online experiment, and their values were estimated at each time point via spatial interpolation from neighboring electrodes. ASR was also applied to suppress transient noise such as eye blinks and muscle activity. The two-minute baseline recording was used as a reference to estimate ASR parameters, and the resulting transformation was applied throughout the online experiment. For offline analyses, the same preprocessing procedure—including filtering, bad channel interpolation, and ASR application—was applied to maintain consistency with the online processing pipeline.

After preprocessing, EEG data were segmented into epochs time-locked to stimulus onset, ranging from −200 ms to 600 ms. Baseline correction was applied by subtracting the mean voltage from the pre-stimulus interval (−200 to 0 ms) from each channel in the epoch. These baseline-corrected epochs were then passed through a pre-trained xDAWN spatial filter. The xDAWN algorithm enhances the discriminability between target and non-target event-related potentials (ERPs) by maximizing the signal-to-signal-plus-noise ratio [29, 31]. In this study, xDAWN filters were trained offline using pre-training data (see Section II. F) and applied without further adaptation during online classification.

### E. Pre-training Dataset

The xDAWN spatial filters and the classifier used in this study were pre-trained on a dataset collected from thirty-seven healthy participants in our previous study [28]. In that experiment, participants performed a four-class P300 BCI task over six sessions, which involved three different stimulus types and two types of mental strategies. Each session consisted of 45 blocks, with each of the four stimuli presented twice per block, resulting in eight sequential presentations per block. This yielded a total of 9,990 target-class samples (45 blocks × 6 sessions × 37 participants) and 29,970 non-target samples, which were three times as numerous as the target class. The protocol and visual paradigm were consistent with those of the current study, making the dataset well-suited for training models in calibration-free online BCI systems.

### F. Classification

The spatially filtered EEG features were input to a deep convolutional neural network (DeepConvNet) [30], which had been trained offline using the pre-training dataset. The DeepConvNet architecture (Fig. 1E) consisted of an initial temporal convolution layer followed by a spatial convolution layer, and then three additional convolutional blocks, each comprising batch normalization, a non-linear activation function (ELU), max pooling, and dropout regularization. This architecture was designed to effectively capture both the temporal dynamics of P300 responses and the discriminative spatial features derived from target and non-target xDAWN components (Fig. 1F). The model produced a binary prediction (target vs. non-target) for each individual stimulus presentation. A final decision for each block was made by selecting the stimulus with the highest posterior probability of being the target among the four presented options. The time to run the pre-trained xDAWN spatial filter per trial was 2.03 milliseconds, and the pre-trained DeepConvNet classifier per trial was 668.43 milliseconds on average (Intel Core i9-12900K CPU, 32 GB DDR4 RAM, NVIDIA GeForce RTX 3060 Ti GPU), supporting real-time BCI system operation.

### G. Offline Analyses

Following the online experiment, several offline analyses were conducted to evaluate system robustness, identify informative EEG channels, and provide empirical guidelines for dataset construction in calibration-free P300 BCIs. First, a channel ablation analysis was performed to assess the contribution of each EEG channel to classification performance. For each channel, the corresponding ERP signal was replaced with zeros, and the classification process—pre-trained xDAWN spatial filters and the DeepConvNet—was re-applied to the entire test dataset. The resulting drop in accuracy was used to quantify the importance of each channel. Second, a data sufficiency simulation was conducted to determine the amount of training data required to build effective pre-trained models. To emulate real-world conditions where BCI datasets are gradually accumulated, we incrementally increased the number of participants (1–37) and blocks per participant (10–270), following the actual data collection order. The lower bound of 10 blocks per participant was chosen to ensure sufficient data for partitioning into training and validation sets, which is necessary for stable DeepConvNet optimization. Data were accumulated in steps of five blocks per participant to balance temporal resolution with computational feasibility. At each iteration, a new xDAWN + deep ConvNet model was trained on the incrementally accumulated data and evaluated using the EEG recordings from the online experiment. To model the trend in decoding accuracy, we fitted an inverse-saturation function of the form:

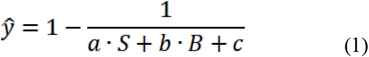

where *ŷ* denotes the predicted classification accuracy, *S* is the number of training subjects, and is the number of blocks per subject. The parameters *a, b*, and were estimated from the data, capturing the respective contributions of subject count, block count, and an intercept term. This inverse saturation function reflects the empirical observation that BCI accuracy asymptotically approaches a ceiling as the amount of training data increases.

Finally, an error trial analysis was conducted to investigate the behavioral and neural correlations of misclassifications. Post-experiment interviews were conducted to collect participants’ subjective accounts of errors, including loss of attention or target uncertainty. Quantitatively, baseline alpha-band power (8–14 Hz) during error trials was compared to that of correct trials to evaluate whether fluctuations in attentional readiness contributed to decoding errors [32–34]. Baseline activity was defined as the pre-stimulus interval (−200 ms to stimulus onset) within each block. In addition, we examined whether the temporal position of the target stimulus within a block influenced error likelihood. To this end, we used a binomial generalized linear mixed model (GLMM), where each trial was treated as an observation. The model included a subject-specific random effect and an intercept-only fixed effect. The intercept’s deviation from the theoretical chance level of 0.25 (equivalent to –1.099 on the logit scale) was assessed using a Z-test. Using GLMM allowed us to assess whether the target position introduced a systematic bias in classification accuracy beyond what would be expected by chance.

### H. Performance Evaluation

BCI performance was assessed using two primary metrics: classification accuracy and ITR. Accuracy was defined as the proportion of test blocks in which the target command was correctly identified by the BCI system. ITR was computed in bits per minute according to the standard BCI formula:

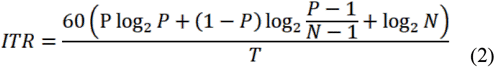

where P denotes the classification accuracy, N represents the number of selectable options (with N=4 in present study), and T refers to the time required for a single selection in seconds, calculated as T=(Stimulus duration)×N×(Number of repetitions per stimulus).

### I. Statistical Analysis

To evaluate the relationship between attentional state and classification performance, we conducted two statistical analyses: a bootstrap test for alpha-band power and a permutation test for temporal error bias. First, to assess prestimulus attentional readiness, we compared alpha power (8–14 Hz) between correct and error trials at each EEG channel. For each of the five posterior and occipital electrodes (P4, P8, O1, Oz, O2), we performed bootstrap resampling (100,000 iterations) from each condition to estimate the distribution of mean differences. Statistical significance was determined by checking whether the 95% confidence interval excluded zero.

Second, to examine whether errors were temporally biased toward particular stimulus positions, we conducted a permutation analysis. For each of 100,000 iterations, we randomly shuffled the target position labels within the set of error trials while preserving the total number of errors. The proportion of errors assigned to each position (1st to 4th) in the actual data was then compared against the resulting null distributions to determine statistical significance. To complement this, we fitted a binomial GLMM for each target position, with the subject as a random effect. The intercept of each model was compared to the theoretical logit value of 0.25 (chance-level selection) using a Z-test.

## III. Results

### A. Online Decoding Performance and Comparison with Calibration-Free Offline Analysis

To evaluate the real-time performance of the proposed calibration-free, single-trial P300-based BCI system, we compared the results from the online experiment with those from the benchmark offline analysis. The offline analysis result was obtained from our previous study on the calibration-free P300-based BCIs using leave-one-subject-out (LOSO) cross-validation [28]. In the offline analysis, EEG data from thrty-seven participants were used to stimulate a subject-independent decoding scenario. Specifically, xDAWN spatial filters and a deep ConvNet classifier were trained using data from thirty-six participants and tested on the remaining participant in each iteration. This process was repeated across all participants. The resulting average classification accuracy was 0.8775 (SD=0.1436), and the corresponding ITR was 52.59 bits/min (SD=21.29). In the online experiment of the present study, the BCI system using the same pre-trained xDAWN and DeepConvNet models yielded the average classification accuracy of 0.8520 (SD=0.1055) and an ITR of 46.41 bits/min (SD=17.77), as shown in Fig. 2. To statistically assess the equivalence of offline and online decoding performance, we applied a Wilcoxon rank-sum test between the online and offline decoding accuracy results. The test revealed no significant difference in accuracy (p=0.1323), indicating that real-time decoding performance was statistically comparable to the offline benchmark. Similarly, no significant difference was observed for ITR (p= 0.1323).

**Fig. 2.**
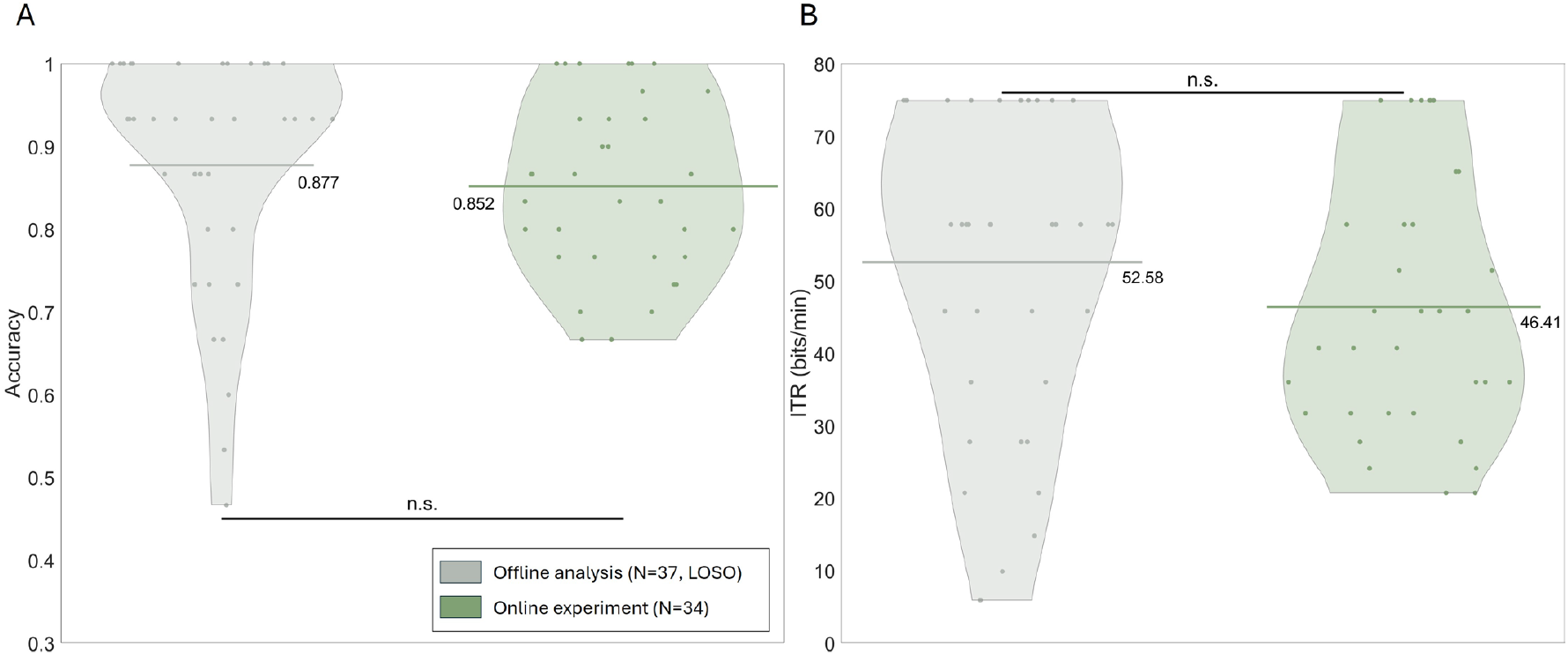
Comparison of classification performance between offline benchmark and online random-order condition. A) Classification accuracy and B) information transfer rate (ITR) are presented for two conditions: calibration-free offline analysis and online decoding under random-order stimulus presentation.

### B. Channel Ablation Analysis for Identifying Channel Contributions

We performed channel ablation analysis to inspect the contribution of individual EEG channels to decoding performance. The results showed that removing five specific channels, including P3, P4, P8, O1, and Oz, led to a statistically significant decrease in decoding accuracy (paired t-test, Bonferroni corrected, p < 0.05) (Fig. 3A). These channels were located in the parietal and occipital regions, which are well known for generating P300 responses. Even among channels that did not meet the corrected significance threshold, ablating electrodes in posterior regions generally resulted in reduced accuracy, suggesting a disproportionately important role in supporting decoding. Fig. 3B shows the topographical distribution of all 31 electrodes, with the five statistically significant channels highlighted. These findings indicate that the proposed P300 BCI system relies primarily on ERPs recorded over parietal and occipital areas and provide data-driven guidance for future optimization of low-density BCI configurations.

**Figure 5.**
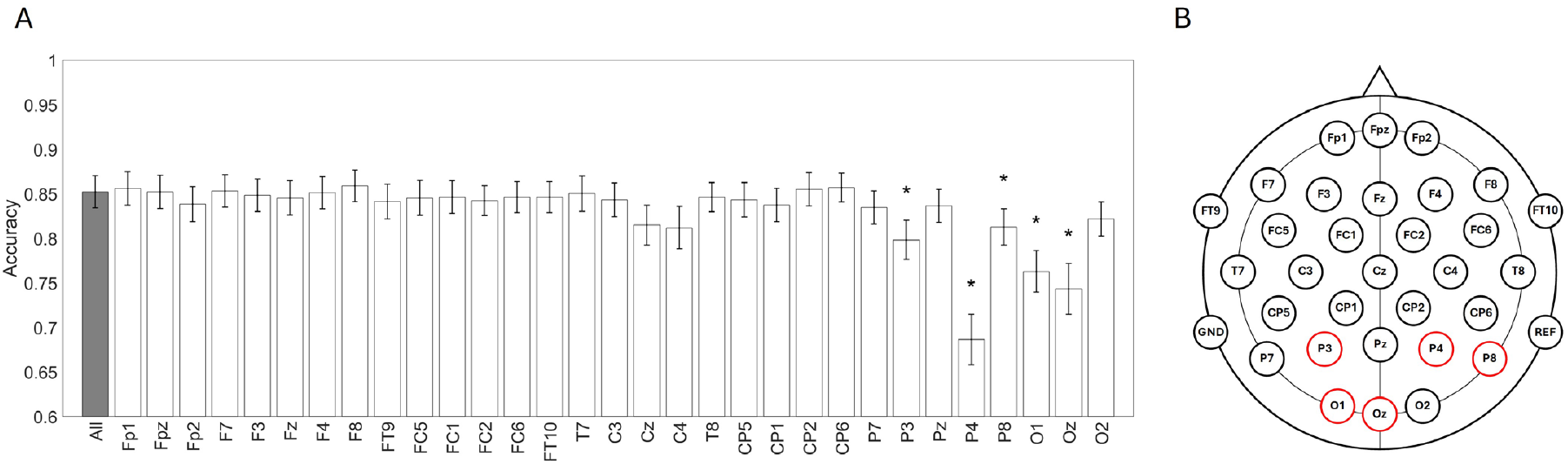
Channel ablation analysis to identify electrode importance. A) Decoding accuracy using the full 31-channel setup (leftmost bar) is compared with accuracy from ablation conditions, where each individual channel was removed (set to zero) one at a time. Error bars indicate standard error. Asterisks mark electrodes whose removal caused a statistically significant performance drop after Bonferroni correction (p < 0.0016). B) Scalp map of the 31 EEG channels used in the study. Channels showing significant impact on accuracy (P3, P4, P8, O1, Oz) are highlighted in color. Results indicate that parietal and occipital electrodes contribute most critically to P300 classification.

### C. Data Sufficiency Simulation for Effective Calibration-Free Training

To determine how much training data is required to construct reliable calibration-free P300-based BCIs, we performed a data sufficiency simulation using a previously collected dataset of 37 participants. We systematically varied the number of participants (1–37) and the number of blocks per participant (10–270) used in training. Fig. 4A illustrates how classification accuracy increased as the size of the training dataset increased. To characterize the observed trend, we fitted the inverse-saturation function to the simulated accuracy data (see Section II. G). The fitted model demonstrated that accuracy increased nonlinearly with dataset size, showing diminishing returns beyond a certain point. The optimal coefficients identified through model fitting were a = 0.1713, b = 0.0063, and c = 1.5525. Nonlinear least squares regression analysis revealed that all three parameters were significantly different from zero (ps < 0.001), indicating that both the number of subjects and the number of blocks per subject significantly contributed to accuracy. Fig. 4B displays the fitted prediction surface and iso-accuracy contours at 0.70, 0.75, 0.80, and 0.85 as representative examples. These contours demonstrate a clear trade-off between the number of participants and the number of blocks per participant. Specifically, when fewer participants are available, more blocks are required to achieve high accuracy. Conversely, with 30 or more participants, models could reach the accuracy of 0.85 using only 10 blocks per subject.

**Fig. 4.**
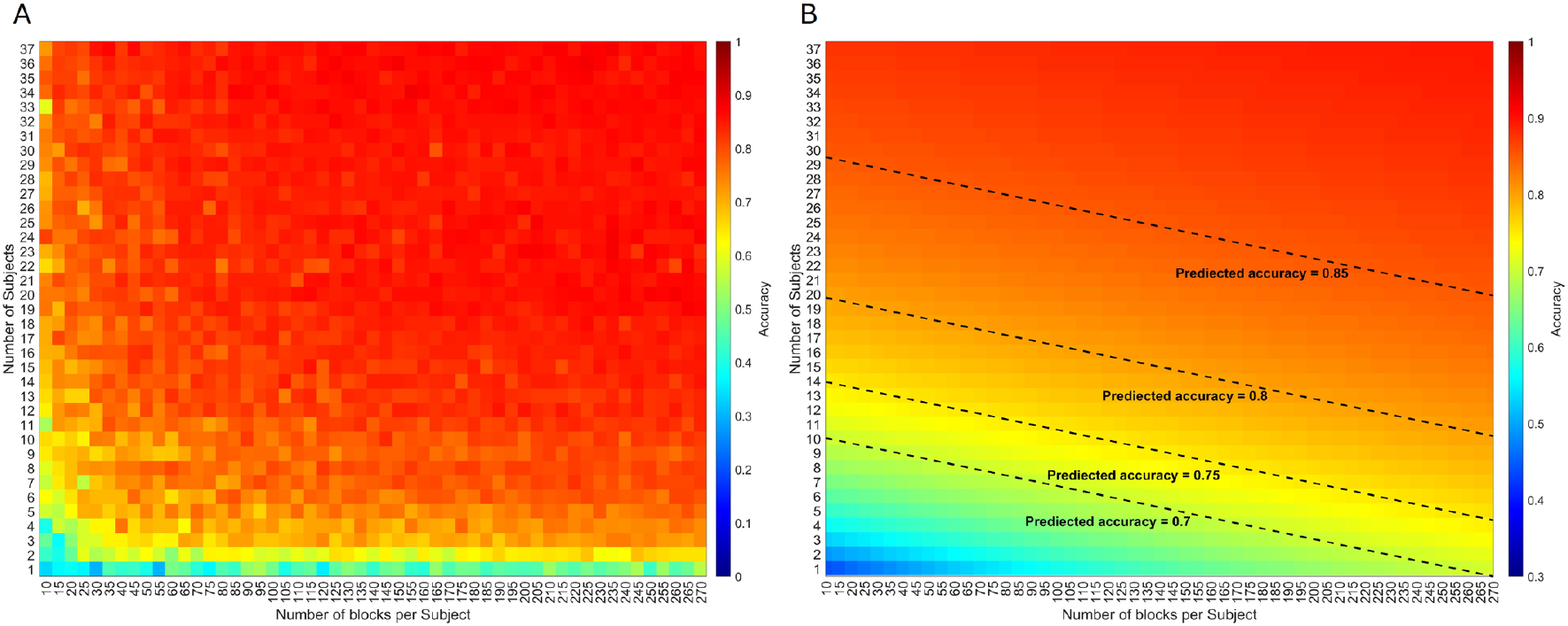
Data sufficiency analysis for calibration-free model training. A) Classification accuracy as a function of the number of training participants and blocks per participant. Accuracy improves with more data, following the actual collection order. B) Predicted accuracy surface from an inverse regression model with overlaid contours indicating target accuracy levels (e.g., 0.70–0.85). Results show that using data from 30 or more subjects requires only ~10 blocks per subject to achieve ≥0.85 accuracy, providing practical guidelines for dataset construction.

### D. Error Trial Analysis

#### 1) Prestimulus Attentional State: Alpha-Band Power

We hypothesized that BCI control errors may be linked to the participant’s attentional state at the beginning of each block. Specifically, we analyzed alpha-band power (8–14 Hz) during the baseline period (−200ms–onset) immediately before the first stimulus appeared. Because the number of error and correct trials varied substantially across participants and were highly imbalanced, we used a bootstrap approach (100,000 iterations) to compare average alpha power between error and correct trials at each EEG channel. The analysis revealed that alpha power was significantly lower in error trials than in correct trials at five posterior and occipital channels: P4 (p = 0.0094), P8 (p = 0.0116), O1 (p = 0.0094), Oz (p = 0.0226), and O2 (p = 0.0267) (Fig 5). This finding suggests that reduced prestimulus alpha activity—often associated with reduced internal attention—corresponds to higher error probability, potentially due to reduced attentional readiness prior to trial onset.

**Fig. 5.**
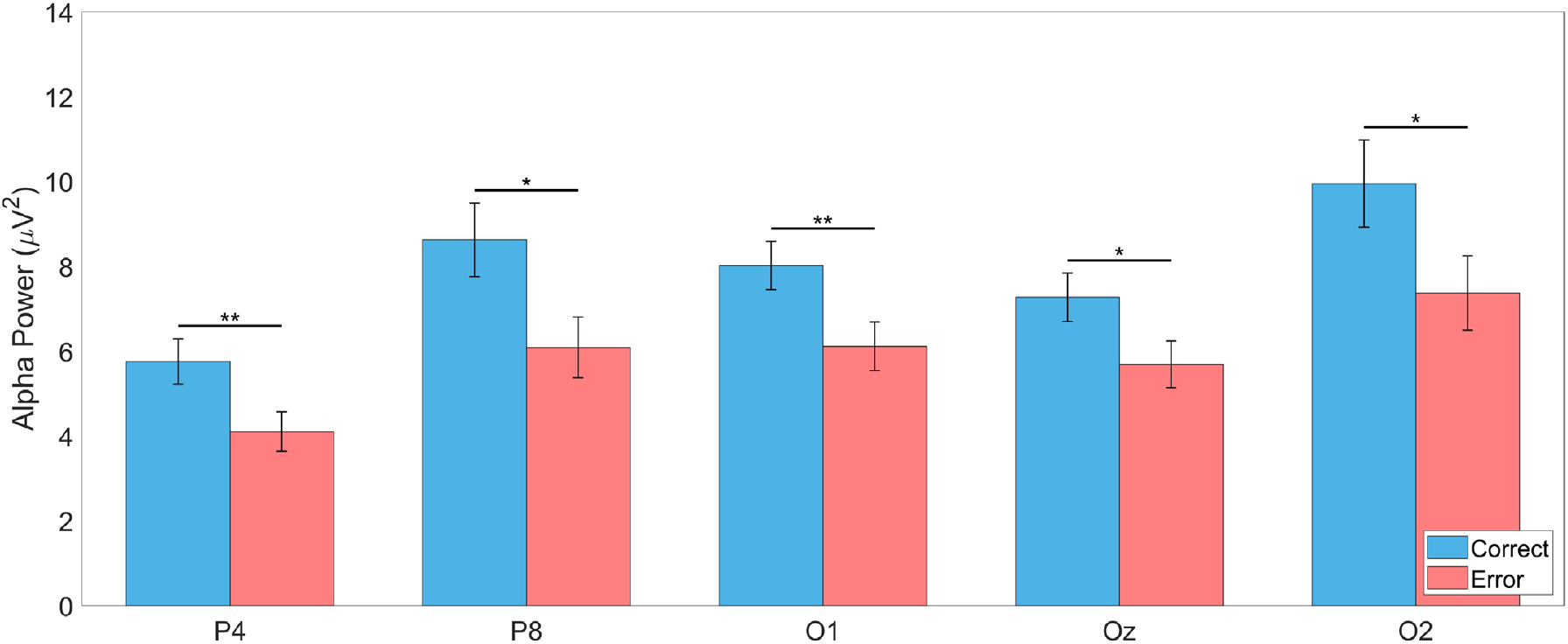
Prestimulus alpha-band power in correct and error trials. Bootstrap-estimated mean and SEM of prestimulus alpha power (−200 to 0 ms) across five posterior electrodes: P4, P8, O1, Oz, and O2. Each pair of bars represents the average alpha power for correct and error trials based on 100,000 bootstrap samples, with error bars indicating standard error of the mean. Across all channels, alpha power was consistently higher in correct trials, suggesting that increased prestimulus alpha activity is associated with enhanced attentional readiness and improved BCI performance. Asterisks indicate channels with statistically significant differences (p < 0.05) where the 95% bootstrap confidence interval of the group difference did not include zero. Asterisks denote significance levels: *p < 0.05, **p < 0.01, ***p < 0.001.

#### 2) Target Presentation Order and Temporal Error Bias

In a single-trial setting, the target stimulus is presented only once. If the participant misses that single target appearance, an error is inevitable. We therefore analyzed whether the position of the target stimulus within the trial sequence (1st to 4th) influenced the likelihood of error. Fig. 6A displays the empirical error rates as a function of target presentation order (1st to 4th). Notably, the error rate was the highest when the target appeared in the 4th position and the lowest in the 2nd. The permutation test revealed that error rates in the 2nd and 3rd positions were significantly lower, and the error rate in the 4th position was significantly higher than expected under the null model (ps < 0.001). To confirm these results, we fitted a binomial GLMM for each target position, treating subject as a random effect. Intercepts were tested against the expected logit for random selection (chance = 0.25). As shown in Fig. 6B, the 4th position had a significantly higher intercept (p = 0.005), and the 2nd position had a significantly lower one (p = 0.002).

**Fig. 6.**
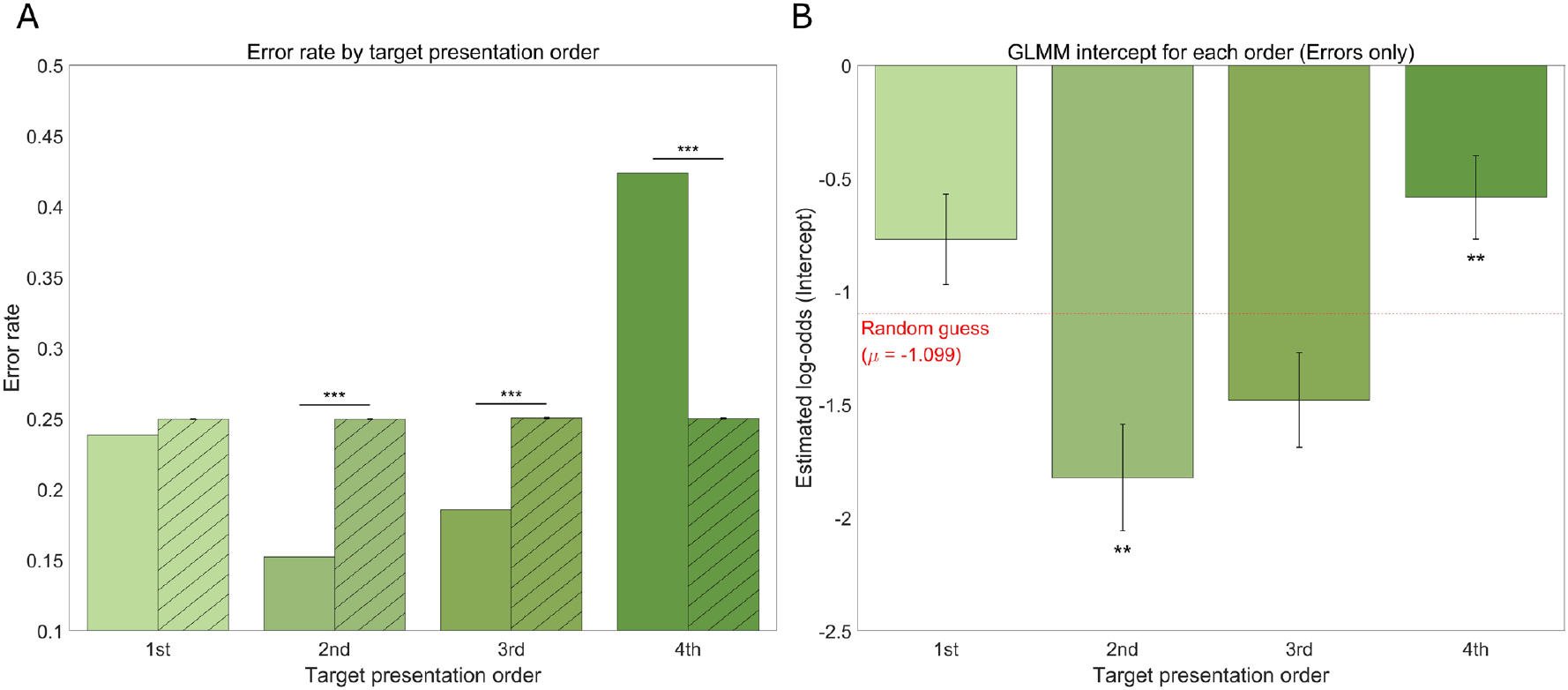
Temporal position effects on classification errors in single-trial BCI. A) Error rates for each target presentation position (1st to 4th) are shown using solid bars for observed data and hatched bars for the null distribution obtained via 100,000 permutations. The 2nd and 3rd positions showed significantly lower error rates than expected by chance, while the 4th position showed a significantly higher error rate. B) Intercepts from a binomial generalized linear mixed model (GLMM) fitted separately for each target position. The red dashed line indicates the theoretical log-odds under random guessing (μ = –1.099). Intercepts for the 2nd and 3rd positions were significantly below chance, indicating improved detection accuracy, while the 4th position approached chance level. Asterisks denote significance levels: *p < 0.05, **p < 0.01, ***p < 0.001.

## IV. Discussion

In this study, we developed and validated a single-trial, calibration-free P300-based BCI system that combines a pre-trained xDAWN spatial filter with DeepConvNet. Unlike previous studies that relied solely on offline validation or required subject-specific calibration, our system demonstrated robust decoding performance in real-time, achieving 85.2% accuracy—comparable to the offline benchmark of 87.8%. The entire processing pipeline was designed to be computationally efficient and robust to noise, enabling low-latency operation without the need for individualized training. Together with a cognitively engaging stimulus paradigm and real-time interaction with IoT devices, these results mark a significant step toward practical, plug-and-play BCI systems that are scalable, user-independent, and ready for real-world deployment.

### A. Comparison to Existing Studies

Compared to prior work on calibration-free P300-based BCIs, our study demonstrates one of the most robust systems to date in terms of decoding performance and participant scale. While several recent studies have pursued subject-independent approaches, few have validated them under a true plug-and-play scenario—i.e., applying a pre-trained model to entirely new users without any subject-specific calibration. Among the previous studies summarized in Table I, only Hu et al. (2024) [35] and our work performed online experiments using a fully pre-trained decoder. In contrast, Gao et al. (2023) incorporated a few calibration trials for adaptation [36], and Huang et al. (2022) conducted only an offline analysis with subject-specific ERP normalization [37]. These differences emphasize that true real-time plug-and-play validation remains rare.

**TABLE 1.**
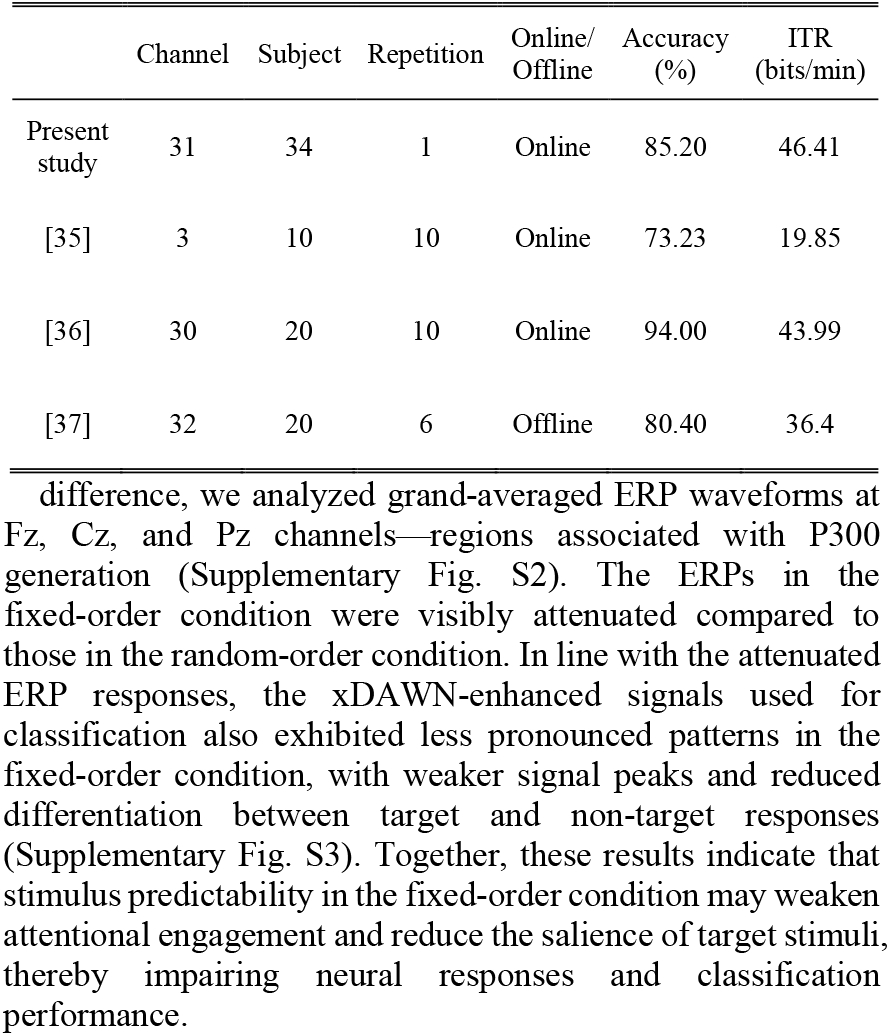
Comparison of classification performance across calibration-free BCI conditions.

|  | Channel | Subject | Repetition | Online/<br>Offline | Accuracy<br>(%) | ITR<br>(bits/min) |
| --- | --- | --- | --- | --- | --- | --- |
| Present study | 31 | 34 | 1 | Online | 85.20 | 46.41 |
| [35] | 3 | 10 | 10 | Online | 73.23 | 19.85 |
| [36] | 30 | 20 | 10 | Online | 94.00 | 43.99 |
| [37] | 32 | 20 | 6 | Offline | 80.40 | 36.4 |

In our system, single-trial decoding was performed without repetition or user-specific tuning yet achieving high accuracy (85.2%)—second among the compared works—and the highest ITR, owing to the single-trial stimulus design. We expect that the accuracy could be further improved if we increase the number of stimulus presentations, as demonstrated in our previous work (Kim et al., 2024). Furthermore, our online BCI system was tested in 34 participants, exceeding the scale of other previous real-time studies. Collectively, our results underscore the practical feasibility and scalability of plug-and-play P300 BCIs.

### B. Implications of The Channel Ablation Analysis

The channel ablation analysis revealed that EEG channels over the parietal and occipital regions (e.g., P3, P4, P8, O1, Oz) made the most substantial contributions to classification performance. This finding aligns with previous neurophysiological findings indicating that the P300 component in visual oddball tasks is most pronounced in these regions [25, 38, 39]. Our results also suggest that a reduced electrode set targeting parieto-occipital regions may preserve decoding performance while simplifying EEG system design.

Unlike previous studies that emphasized subject-specific optimization of EEG channel selection [40–42], our approach evaluates channel importance in a cross-subject, pre-trained computational framework. This generalizable approach aims to identify a consistent set of informative channels applicable across users, providing a foundation for constructing calibration-free BCI systems with fixed, low-density electrode layouts. Such configurations are critical for improving usability and enabling plug-and-play deployment in daily life.

### C. Evaluation of Dataset Sufficiency and Model Generalization

The data sufficiency simulation showed the relationship between the training dataset size and expected classification performance. The inverse regression model predicted that with 30 or more participants, as few as 10 blocks per subject are sufficient to achieve an average accuracy above 0.85. Notably, our result indicates that collecting data from a large number of people would be more effective than collecting many data samples from a small number of people. However, collecting large-scale pre-training datasets is often nontrivial, especially under custom experimental paradigms such as ours. Moreover, public EEG datasets were not directly compatible due to differences in stimulus design and task structure [43].

To further examine the validity of the model prediction, we conducted an additional bi-directional evaluation: we used the online experimental data from 34 participants (random-order condition only) as the training set and applied the resulting pre-trained model to the original 37-participant dataset, which had previously been used to train the system. In other words, the training and test sets were swapped. This analysis yielded an average decoding accuracy of 0.8198 (SD = 0.1490) on the 37-subject test set. The predicted accuracy from our regression model for the same training size was 0.8547, indicating a moderate deviation between predicted and observed performance. While the model slightly overestimated accuracy, the actual result remains within a reasonable margin and validates the predictive capability of the model.

### D. Target Presentation Order and Error Bias

The analysis of error trials revealed that target stimuli presented later in the trial sequence—particularly in the fourth position—were more likely to result in classification errors. Post-experiment interviews with participants provided qualitative support for this observation. Several participants noted that, due to the fixed temporal rhythm of the stimulus sequence, they could anticipate the timing of the fourth stimulus. As a result, when the target was positioned last, the stimulus no longer appeared as temporally rare or unexpected—key conditions for eliciting a robust P300 response in an oddball paradigm. This predictability may have reduced the salience of the target stimulus and diminished attentional engagement at the critical moment. Interestingly, the error rate for trials in which the target appeared in the second position was significantly lower than expected by chance. One possible explanation is that, at this early stage of the stimulus sequence, participants were still actively maintaining attention and had not yet begun to mentally disengage or form premature expectations about the target’s timing. Additionally, the second stimulus may benefit from being early enough to retain novelty while not being so early as to catch participants off-guard, as the first stimulus might sometimes do. This result suggests that the temporal dynamics of stimulus presentation—beyond mere randomness—can meaningfully influence decoding performance.

These behavioral findings align with our neural analysis of prestimulus alpha-band activity. While previous studies have associated alpha desynchronization (i.e., reduced alpha power) with increased visual attention during stimulus processing [25, 44–47], our study focused on the preparatory period prior to stimulus onset. During this anticipatory phase, higher alpha power has been interpreted as a marker of top-down inhibition of irrelevant neural processing [32–34]. We found that alpha power preceding error trials was significantly lower than that preceding correct trials, particularly at posterior sites. This may indicate insufficient inhibition during preparatory attention, possibly due to participants’ momentary lapses in vigilance. Taken together, these findings suggest that not only the randomness of target location but also the temporal unpredictability of its appearance is critical for maintaining attentional readiness and eliciting robust P300 responses. Practically, it underscores the importance of optimizing stimulus timing and incorporating attentional state monitoring (e.g., alpha power) into future adaptive BCI systems.

### E. ERP and Feature Representation Differences between Fixed and Random Conditions

In addition to the main random-order stimulus presentation, we performed an additional online experiment under a fixed-order stimulus presentation condition. This design choice was motivated by a practical concern specific to single-trial BCIs: in real-time use, missing a single target presentation leads directly to classification failure, since the stimulus is not repeated. We hypothesized that increasing the temporal predictability of stimulus appearance—by presenting stimuli in a fixed spatial order (e.g., clockwise)—might help users better anticipate the timing of their target and thus avoid attentional lapses. To evaluate this possibility, we tested an additional group of 16 participants (mean age = 22.44 years, SD = 3.30) under the fixed-order condition.

Although both experimental conditions in this study followed a four-class oddball paradigm, we observed notable differences in neural responses and classification performance between the fixed-order and random-order settings. In the fixed-order condition, where the spatial presentation sequence of stimuli was consistent across blocks (Fig. 1D), the average classification accuracy was 0.7146 (SD = 0.1398), and the corresponding ITR was 28.40 bits/min (SD = 16.82). In contrast, the random-order condition yielded significantly higher accuracy as reported in Section III. A (Supplementary Fig. S1A) (0.8520, SD = 0.1055) and ITR (Supplementary Fig. S1B) (46.41 bits/min, SD = 17.77), along with lower inter-subject variability. A Kruskal–Wallis test followed by post hoc Tukey–Kramer analysis confirmed that the fixed-order performance was significantly lower than the random-order condition (p=0.0124). These findings suggest that temporal unpredictability preserves attention and maintains online BCI performance across users.

To further explore the neural basis of this performance difference, we analyzed grand-averaged ERP waveforms at Fz, Cz, and Pz channels—regions associated with P300 generation (Supplementary Fig. S2). The ERPs in the fixed-order condition were visibly attenuated compared to those in the random-order condition. In line with the attenuated ERP responses, the xDAWN-enhanced signals used for classification also exhibited less pronounced patterns in the fixed-order condition, with weaker signal peaks and reduced differentiation between target and non-target responses (Supplementary Fig. S3). Together, these results indicate that stimulus predictability in the fixed-order condition may weaken attentional engagement and reduce the salience of target stimuli, thereby impairing neural responses and classification performance.

### F. General Implications and Future Directions

Overall, our results demonstrate that calibration-free, single-trial P300 BCI systems can operate effectively in real-time. The ability to train decoders on pre-existing data and decode new users’ intentions without subject-specific calibration represents a meaningful advance in BCI usability. In addition, our error trial analysis indicated that prestimulus alpha-band power—reflecting preparatory attentional state—was significantly lower prior to incorrect classifications. This suggests that real-time monitoring of alpha power may enable online system adjustments (e.g., prompting the user to refocus or delaying stimulus onset) to mitigate attentional lapses. Furthermore, our comparison of fixed and random stimulus sequences revealed that temporal unpredictability enhances attentional engagement and neural discriminability. Together, these insights point to the future work of implementing adaptive timing mechanisms, such as randomized inter-stimulus intervals, to prevent user fatigue and maintain optimal attentional states throughout BCI operation. Future work should prioritize the scaling of pre-training datasets, refinement of low-channel configurations, and incorporation of real-time system adjustments. Our results contribute to the growing body of evidence that practical, generalizable BCIs are within reach and lay a foundation for their integration into everyday applications.

## V. Conclusion

This study presents the real-time validation of a calibration-free, single-trial P300-based BCI system operating under ecologically realistic conditions. By leveraging a pre-trained xDAWN spatial filter and a deep convolutional neural network—trained exclusively on data from independent users—we successfully enabled plug-and-play BCI use without any subject-specific calibration. The online system achieved classification performance comparable to that of offline benchmarks. Key design elements contributing to this success included the use of randomized stimulus sequences to maintain attentional novelty, minimal yet effective signal preprocessing, and a compact deep learning architecture suitable for real-time deployment. In addition to the real-time results, our offline analyses established quantitative guidelines for training data sufficiency and identified key electrode locations for low-density implementation. Together, these findings validate the feasibility of real-time, calibration-free P300 BCI systems and offer practical strategies for improving robustness and scalability. The proposed framework provides a strong foundation for future research aiming to develop fully deployable, user-adaptive plug-and-play BCIs that extend beyond laboratory settings into real-world use.

