## Supplemental Figure S1, Supplemental Figure S2, Supplemental Figure S3, and Supplemental video for "A Plug-and-Play P300-Based BCI with Calibration-Free Application"

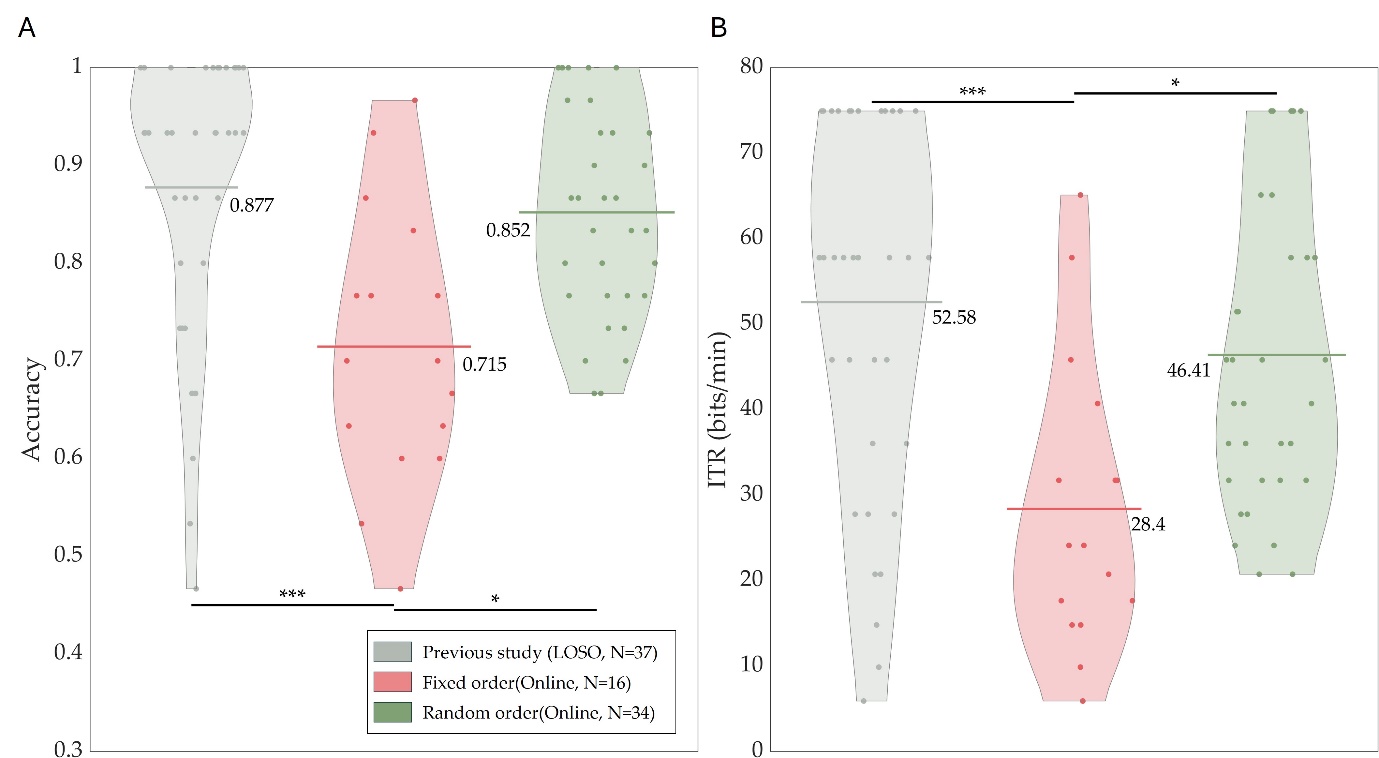

**Fig. S 1. Comparison of classification performance between offline benchmark and online random-order condition.**
A) Classification accuracy and B) information transfer rate (ITR) are presented for two conditions: calibration-free offline analysis and online decoding under random-order stimulus presentation. Asterisks denote significance levels: *p < 0.05, **p < 0.01, ***p < 0.001.

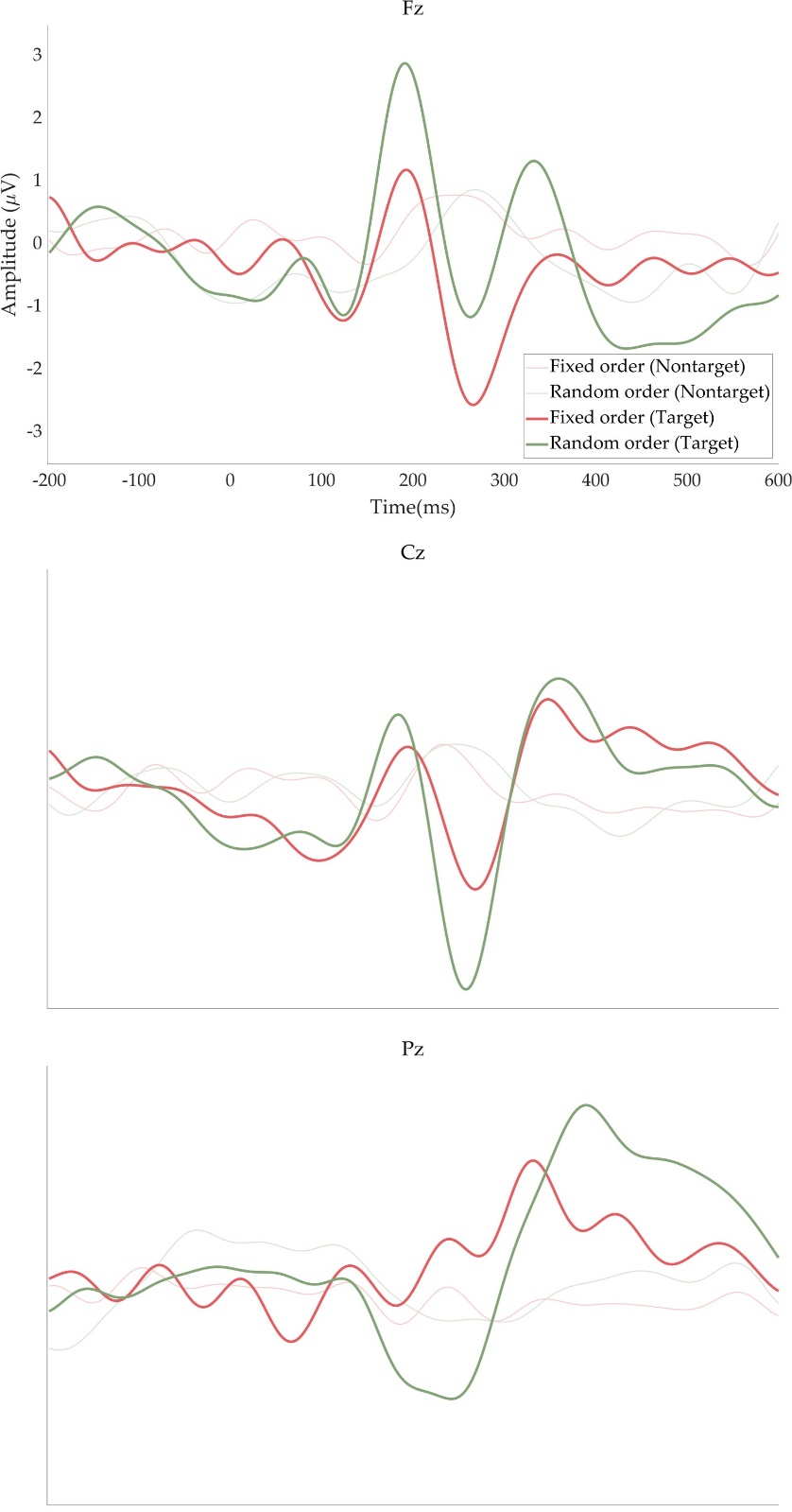

**Fig. S 2. Grand-averaged ERP waveforms from representative electrodes in fixed and random stimulus order conditions.** ERP signals recorded at Fz, Cz, and Pz are shown separately for the fixed-order (left) and random-order (right) online conditions. Each plot displays the average across all participants. The thick line represents the ERP for target stimuli, while the thin line corresponds to non-target stimuli. Clearer and more pronounced P300 components are observed in the random-order condition across all electrodes, suggesting stronger attentional engagement and improved target–non-target discrimination when stimulus unpredictability is maintained.

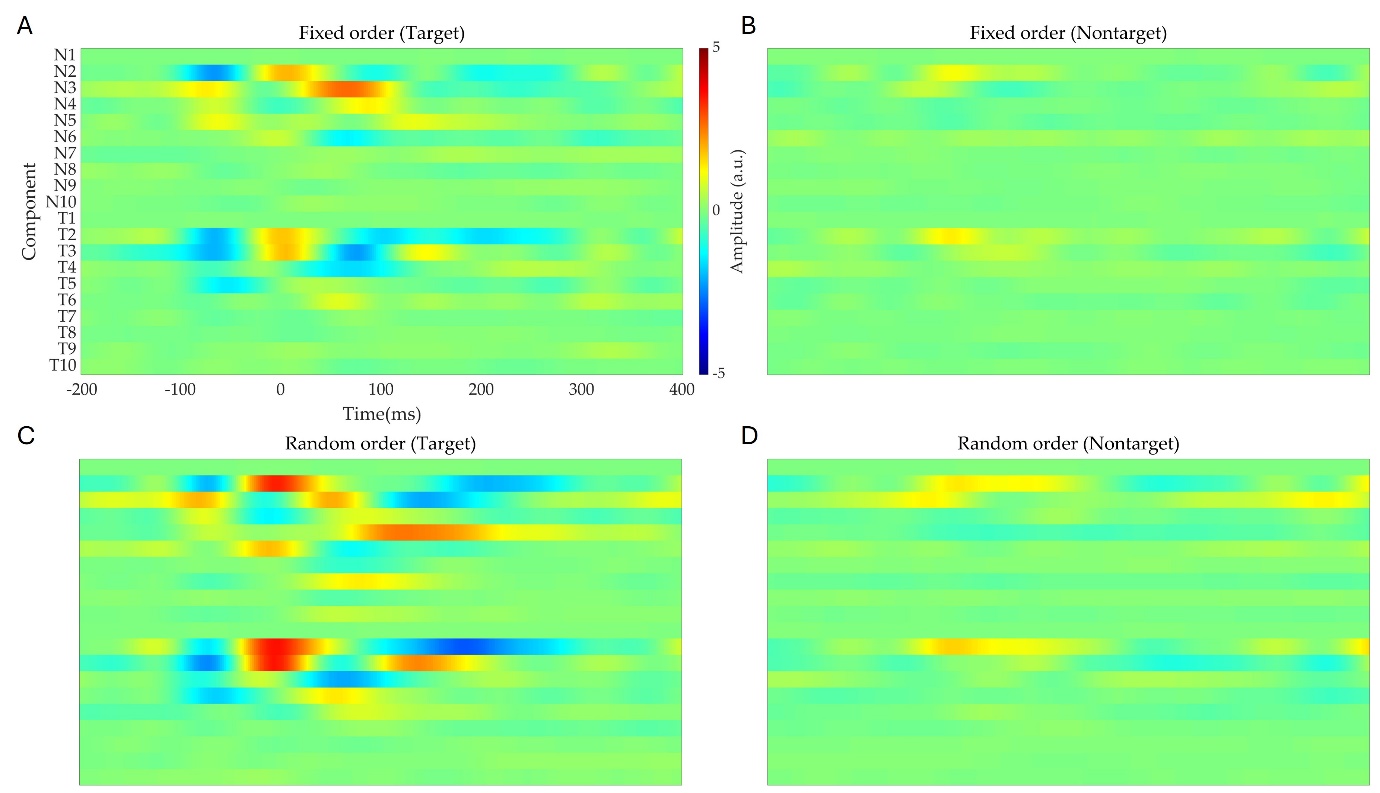

**Fig. S 3. xDAWN-filtered signal representations used as input to the ConvNet classifier.** The plots show the average input features across participants after applying the xDAWN spatial filter (with 10 components) for each condition and label type. A) Fixed-order condition, target trials (T1–T10); B) Fixed-order condition, non-target trials (N1–N10); C) Random-order condition, target trials (T1–T10); D) Random-order condition, non-target trials (N1–N10).

**Supplementary video 1**

<https://www.youtube.com/watch?v=AzOJIez4XjU>
